# Robustness of STDP to spike timing jitter

**DOI:** 10.1101/259648

**Authors:** Yihui Cui, Ilya Prokin, Alexandre Mendes, Hugues Berry, Laurent Venance

## Abstract

In Hebbian plasticity, neural circuits adjust their synaptic weights depending on patterned firing of action potential on either side of the synapse. Spike-timing-dependent plasticity (STDP) is an experimental implementation of Hebb’s postulate that relies on the precise order and the millisecond timing of the paired activities in pre- and postsynaptic neurons. In recent years, STDP has attracted considerable attention in computational and experimental neurosciences. However, canonical STDP is assessed with deterministic (constant) spike timings and time intervals between successive pairings, thus exhibiting a regularity that strongly differs from the biological variability. Hence, the emergence of STDP from noisy neural activity patterns as expected in *in vivo*-like firing remains unresolved. Here, we used noisy STDP stimulations where the spike timing and/or the interval between successive pairings were jittered. We explored with a combination of experimental neurophysiology and mathematical modeling, the impact of jittering on three distinct forms of STDP at corticostriatal synapses: NMDAR-mediated tLTP, endocannabinoid-mediated tLTD and endocannabinoid-mediated tLTP. As the main result, we found a differential sensitivity to jittered spike timing: NMDAR-tLTP was highly fragile whereas endocannabinoid-plasticity (tLTD and tLTP) appeared more resistant. Moreover, when the frequency or the number of pairings was increased, NMDAR-tLTP became more robust and could be expressed despite strong jittering of the spike timing. Taken together, our results identify endocannabinoid-mediated plasticity as a robust form of STDP while the sensitivity to jitter of NMDAR-tLTP varies with activity frequency. This provides new insights into the mechanisms at play during the different phases of learning and memory and the emergence of Hebbian plasticity in *in vivo*-like firing.

## INTRODUCTION

Learning and memory are mainly underlain by long-term synaptic efficacy changes between neurons [1,2] that can be assessed with a synaptic Hebbian paradigm such as spike timing-dependent plasticity (STDP). In STDP, the occurrence of timing-dependent-long-term potentiation (tLTP) or -depression (tLTD) relies on the precise (milliseconds) relative timing of paired pre- and postsynaptic action potentials [3,4]. Since its discovery, STDP has been attracting a lot of interest in computational and experimental neuroscience because it relies on spike correlation and has emerged as a candidate mechanism for experience-dependent changes in the neural circuit, including map plasticity [4,5,6]. STDP depends on 1) the relative timing between pre- and postsynaptic spikes (Δ*t*_STDP_) [4], 2) the number of paired spikes (*N*_pairings_) [7,8,9], 3) the frequency of the paired spikes (*F*_pairings_) [7,10] and 4) membrane depolarization [10,11].

STDP is classically investigated experimentally using regular repetitions of the same spike timing and fixed intervals between successive paired stimulations. A typical experimental protocol consists in pairing pre- and postsynaptic stimulations with a fixed Δ*t*_STDP_ (ranging from −30 to +30 ms for plasticity induction). The pairings are then repeated between 15 and 200 times with a constant time interval between successive pairings (typically between 0.1 and 5 Hz). The impact on STDP of the frequency [7,10,12] and number [7,8,9,12] of pairings has been demonstrated. However, in those studies, the spike timing and the time interval between successive pairings were deterministic.

Regular stimulation paradigms produce patterns of activity that are likely to differ from the variability expected in *in vivo*-like firing. A recent theoretical study has started to explore more naturalistic stimulations [13]. However, whether STDP emergence and maintenance is robust against biological variability remains to be investigated. To address this question, it is important to take into account various forms of STDP, involving distinct intracellular signal transduction pathways, i.e. NMDAR-, mGluR- or endocannabinoid-mediated STDP [4,14]. It is thus expected that those various STDP forms might exhibit different robustness to spike train variability.

In the present study, we evoke STDP with noisy spike timings Δ*t*_STDP_ and analyze the impact of this jittering of the spike timing on the expression of three forms of STDP, both in electrophysiological recordings and in a mathematical model of the underlying signaling pathways. We explore STDP at corticostriatal synapses between cortical pyramidal neurons (somatosensory cortex layer 5) and medium-sized spiny neurons (MSNs) of the dorso-lateral striatum. Corticostriatal STDP has been shown to exhibit three forms of STDP depending on *N*_pairings_: an NMDAR-mediated tLTP and an endocannabinoid-mediated tLTD for *N*_pairings_~100 pairings [15,16,17,18] and an endocannabinoid (eCB)-mediated tLTP STDP for *N*_pairings_~5–10 [8,9].

Combining electrophysiological experiments with the mathematical model, we show that NMDAR- and endocannabinoid**-**STDP forms are not equal with respect to jittering of the spike timing and the pairing frequency: at low frequency (1Hz), eCB-mediated plasticity (eCB-tLTD and eCB-tLTP) appears robust to jittering whereas NMDAR-tLTP is very fragile, disappearing with even small jitter. However, increasing the number of pairings or the average frequency greatly improves NMDAR-tLTP robustness. Our results further suggest that the robustness of NMDAR-tLTP to jitter is also strengthened by the irregularity of the spike-train stimulations. Taken together, our results show that the robustness of STDP depends both on the STDP form under consideration and on the properties of the stimulations (number of pairings, frequency and variability), thus suggesting that the emergence and expression of STDP forms *in vivo* depends on the characteristics of the pre- and postsynaptic inputs.

## METHODS

### Animals and brain slice preparation

OFA rats P_23-32_ (Charles River, L’Arbresle, France) were used for brain slice electrophysiology. All experiments were performed in accordance with the guidelines of the local animal welfare committee (Center for Interdisciplinary Research in Biology Ethics Committee) and the EU (directive 2010/63/EU). Every precaution was taken to minimize stress and the number of animals used in each series of experiments. Animals were housed in standard 12-hour light/dark cycles and food and water were available *ad libitum*.

Horizontal brain slices containing the somatosensory cortical area and the corresponding corticostriatal projection field were prepared as previously described [19]. Corticostriatal connections (between somatosensory cortex layer 5 and the dorso-lateral striatum) are preserved in the horizontal plane. Brain slices (330 µm-thick) were prepared with a vibrating blade microtome (VT1200S, Leica Microsystems, Nussloch, Germany). Brains were sliced in an ice-cold cutting solution (125 mM NaCl, 2.5 mM KCl, 25 mM glucose 25 mM NaHCO_3_, 1.25 mM NaH_2_PO_4_, 2 mM CaCl_2_, 1 mM MgCl_2_, 1 mM pyruvic acid) through which 95% O_2_/5% CO_2_ was bubbled. The slices were transferred to the same solution at 34°C for one hour and then to room temperature.

### Patch-clamp recordings

Patch-clamp recordings were performed as previously described [9,18]. Briefly, for whole-cell recordings in borosilicate glass pipettes of 5-7 MΩ resistance were filled with (in mM): 122 K-gluconate, 13 KCl, 10 HEPES, 10 phosphocreatine, 4 Mg-ATP, 0.3 Na-GTP, 0.3 EGTA (adjusted to pH 7.35 with KOH). The composition of the extracellular solution was (mM): 125 NaCl, 2.5 KCl, 25 glucose, 25 NaHCO_3_, 1.25 NaH_2_PO_4_, 2 CaCl_2_, 1 MgCl_2_, 10 µM pyruvic acid bubbled with 95% O_2_ and 5% CO_2_. Signals were amplified using with EPC9-2 and EPC10-2 amplifiers (HEKA Elektronik, Lambrecht, Germany). All recordings were performed at 34°C, using a temperature control system (Bath-controller V, Luigs&Neumann, Ratingen, Germany) and slices were continuously superfused with extracellular solution, at a rate of 2 ml/min. Slices were visualized under Olympus BX51WI microscopes (Olympus, Rungis, France), with a 4×/0.13 objective for the placement of the stimulating electrode and a 40×/0.80 water-immersion objective for the localization of cells for whole-cell recordings. Current-clamp recordings were filtered at 2.5 kHz and sampled at 5 kHz and voltage-clamp recordings were filtered at 5 kHz and sampled at 10 kHz, with the Patchmaster v2×32 program (HEKA Elektronik).

### Spike timing-dependent plasticity protocols: regular and jittered patterns

Electrical stimulations were performed with a concentric bipolar electrode (Phymep, Paris, France) placed in layer 5 of the somatosensory cortex. Electrical stimulations were monophasic, at constant current (ISO-Flex stimulator, AMPI, Jerusalem, Israel). Cortical stimulations evoked glutamatergic excitatory postsynaptic currents (EPSCs) (inhibited by CNQX 10 µM and D-AP5 50 µM, n=6) and not significantly affected by GABAergic events; Indeed, blocking GABA_A_Rs with picrotoxin (50 µM) did not significantly affect EPSC amplitude at corticostriatal synapses (123±31 pA before and 118±26 pA after picrotoxin, *p*=0.500, n=5). Repetitive control stimuli were applied at 0.1 Hz. Currents were adjusted to evoke 50-200 pA EPSCs. STDP protocols consisted of pairings of pre- and postsynaptic stimulations (at 1 Hz) separated by a specific time interval (Δ*t*_STDP_). The paired stimulations were applied at 1Hz throughout the study except in Figure 6a1 and 6a2 in which 3 Hz stimulations were tested. Presynaptic stimulations corresponded to cortical stimulations and the postsynaptic stimulation of an action potential evoked by a depolarizing current step (30 ms duration) in MSNs. Δ*t*_STDP_ <0 ms for post-pre pairings, and Δ*t*_STDP_ >0 ms for pre-post pairings. Recordings on neurons were made over a period of 10 minutes at baseline, and for at least 60 minutes after the STDP protocols. We individually measured and averaged the amplitude of 60 successive EPSCs from both baseline and 45-55 minutes after STDP protocol, in which the latter was normalized by the former to calculate long-term synaptic efficacy changes. Neuron recordings were made in voltage-clamp mode during baseline and for the 60 minutes of recording after the STDP protocol, and in current-clamp mode during STDP protocol. Experiments were excluded if input resistance (Ri), measured every 10 sec all along the experiment, varied by more than 20%. After recording of 10 min control baseline, drugs were applied in the bath. A new baseline with drugs was recorded after a time lapse of 10 min (to allow the drug to be fully perfused) for 10 min before the STDP protocol. Drugs were present until the end of the recording. All chemicals were purchased from Tocris (Ellisville, MO, USA), except for picrotoxin (Sigma). DL-2-amino-5-phosphono-pentanoic acid (D-AP5; 50 µM) and 6-cyano-7-nitroquinoxaline-2,3-dione (CNQX; 10 µM) were dissolved directly in the extracellular solution and bath applied. N-(piperidin-1-yl)-5-(4-iodophenyl)-1-(2,4-dichlorophenyl)-4-methyl-1H-pyrazole-3-carboxamide (AM251; 3 µM) and picrotoxin (50 µM) were dissolved in ethanol and added to the external solution, such that the final concentration of ethanol was 0.01-0.1%.

For the jittered Δ*t*_STDP_ patterns, we used the following algorithm (cf Protocol 2, below): the time for each presynaptic stimulation was deterministically set by the pairing frequency: 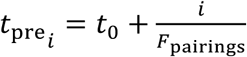 where t_0_ is a constant, usually set to (2*F*_pairings_)^−1^. The postsynaptic times were then chosen randomly according to 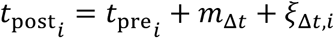 where *m*_Δ*t*_ is the average spike timing and *ξ*_Δ*t*_ is a random variable with mean 0 and variance σ_Δ*t*_^2^.

### Electrophysiological data analysis

Off-line analysis was performed with Fitmaster (Heka Elektronik) and Igor-Pro 6.0.3 (Wavemetrics, Lake Oswego, OR, USA). Statistical analysis was performed with Prism 5.02 software (San Diego, CA, USA). In all cases “n” refers to an experiment on a single cell from a single slice. All results are expressed as mean ± SEM in the text and as mean ± SD in the figures. Statistical significance was assessed in unpaired *t* tests, one way Anova, or in one-sample *t* tests, as appropriate, using the indicated significance threshold (*p*).

### Mathematical model

#### Jittered STDP protocols

A STDP protocol specifies the inter-pairing interval 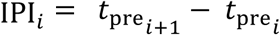 and the spike timing 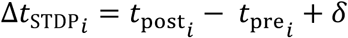. Here,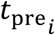 and 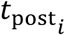 are the times at which the *i*^th^ pre- and postsynaptic stimulations, respectively, are delivered, and δ is the time elapsed between the onset of the postsynaptic step current and the action potential it triggers (around 3 ms in MSNs). In the present study, we introduce stochasticity of the spike timing by adding a random jitter *ξ*_Δ*t*_ to the spike timing: Δ*t*_STDP_ = *m*_Δ*t*_ + *ξ*_Δ*t*_, were *m*_Δ*t*_ is the average spike timing and *ξ*_Δ*t*_ is a random variable whose distribution is given by the STDP protocol. Here, we explored five STDP protocols:

1. *Protocol 0* consisted of the canonical STDP with no jittering, i.e. 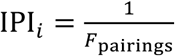 and *ξ*_Δ*t*_ = 0.
2. *Protocol 1* consisted of deterministic IPIs and uniformly distributed spike timings (Fig. 2A1), i.e. 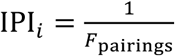, ∀*i*, where *F*_pairings_ is the pairing frequency and the probability distribution function of the jitter is given by

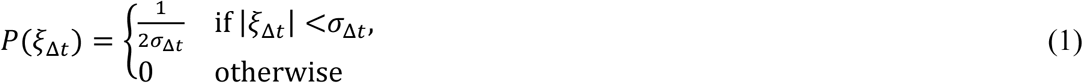

In eq. (1), σ_Δ*t*_ defines the maximal jitter amplitude i.e. the level of noise in the protocol. To simulate *Protocol 1*, we first set each presynaptic stimulation time using 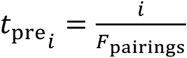. Postsynaptic times were then fixed by 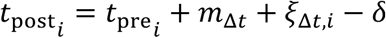.
3. *Protocol 2* (Fig. 2C1) shared the same definition as Protocol 1, except that the jitter followed a normal distribution with zero mean and variance σ_Δt_^2^: P(*ξ*_Δ*t*_) = 𝒩(0, σΔt^2^).
4. For *Protocol 3* (Fig. 2D1), we used a triangle distribution of the jitter by adding uniformly distributed jitter to both presynaptic and postsynaptic times. We set 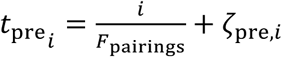 and 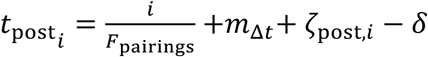, where *ζ*_pre_ and ζ_post_ are i.i.d. random variables with the uniform distribution of eq. (1). The resulting spike timing is *Δt*_STDP_=*m*_Δ*t*_ + *ξ*_Δ*t*_ where *ξ*_Δ*t*_ = ζ_post_ – ζ_pre_ has a triangular distribution. Note that in this case, one still has 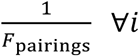 as long as 2|*m*_Δ*t*_|*F*_pairings_ ≪ 1, which can safely be assumed here since we used |m_Δ*t*_| < 50 ms and F_pairings_ < 2 Hz.
5. *Protocol 4* includes both a stochastic spike timing and stochastic IPIs (Fig. 2E1). We first sampled *N*_pairings_ IPIs according to an exponential distribution with rate *λ* and refractory period *τ*_*r*_: P(IPI) = Θ (IPI – *τ*_*r*_) *λ*exp(-*λ*(IPI – *τ*_*r*_)), where Θ(*x*) is the Heaviside function Θ(*x*)=1if *x* ≥ 0; 0 otherwise. The spike trains defined with protocol 4 are thus Poisson process. We then used these IPIs to fix the stimulation times, adding a triangularly distributed jitter as in Protocol 3 above: 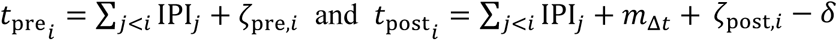. To keep the average stimulation frequency at 1/*F*_pairings_, we constrained the value of the rate *λ* as *λ* = (1/*F*_pairings_ –*τ*_*r*_)^−1^.

#### Stimulations

A detailed account of our mathematical model can be found in S1 Text. We modeled glutamate concentration in the synaptic cleft, *G*(*t*), as a train of exponentially-decaying impulses triggered by presynaptic stimuli at time 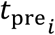:

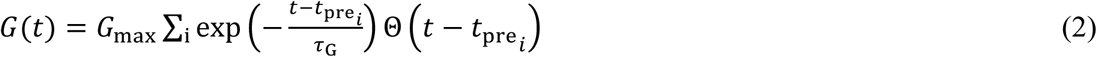

where *G*_max_ is the peak glutamate concentrations and *τ*_G_ is the glutamate clearance rate. We modeled the electrical response to these stimulations in a postsynaptic element considered as a single isopotential compartment with AMPAR, NMDAR, VSCC and TRPV1 conductances

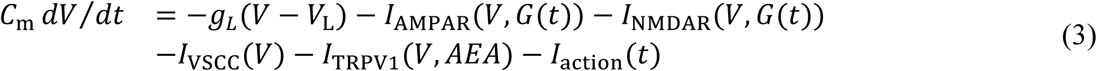

where *V* is membrane potential; *g*_L_ and *V*_*L*_ are leak conductance and reversal potential respectively; *I*_AMPAR_, *I*_NMDAR_, *I*_VSCC_ and *I*_TRPVI_ are currents through AMPAR, NMDAR, L-type VSCC (v1.3) and TRPV1 respectively and AEA stands for anandamide (see below). *I*_action_ *is* the action current resulting from the postsynaptic stimulation (backpropagating action potential on top of a 30 ms depolarization). Details about the analytical expressions of these currents are given in S1 Text.

#### Biochemical signaling

We modeled the kinetics of the biochemical pathways activated by the above electrical stimulations using the model of Cui et al., 2016 [9]. We give below a quick overview of this model and refer to S1 Text for the details.

Free cytosolic calcium is one of the main signaling actors in the model. To model its dynamics, we assumed calcium can be transferred from/to two main sources: (*i*) extracellular calcium, via the plasma membrane channels of eq. (3) above and (*ii*) the endoplasmic reticulum (ER), via the IP3-dependent Calcium-Induced Calcium Release (CICR) system. Hence, the concentration of free cytosolic calcium *C* was computed according to

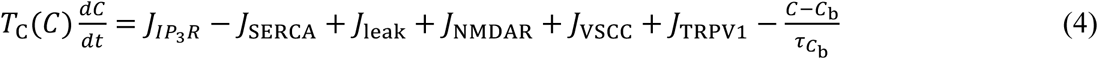

where *J*_IP3R_, *J*_SERCA_, *J*_leak_ are fluxes from and to the ER in the CICR system (see [9]), *J*_NMDAR_ +*J*_VSCC_ +*J*_TRPVI_ are the calcium fluxes from the plasma membrane channels (eq. 3), computed as *J*_x_ = *ξ*_*x*_ ⋅ *I*_*x*_ were the *ξ*_*x*_ are constants, x ∈ {NMDAR,VSCC,TRPV1}. *C*_b_ is the basal cytosolic calcium level resulting from equilibration with calcium diffusion out of the cell and 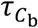 is the corresponding time scale; *T*_*c*_ is a time scaling factor resulting from the presence of endogenous calcium buffers.

In the model, the calcium transients generated by eq. (4) activated a network of biochemical pathways that collectively set the synaptic weight. Hence, in this model, the synaptic weight is entirely fixed by the underlying biochemical signaling network [9]. More precisely, we assumed that the total synaptic weight is the product of a pre- and a postsynaptic contribution *W*_total_ = *W*_pre_*W*_post_. Postsynaptic plasticity was based on the activation by calcium of calmodulin and CaMKII and the regulation of this system by PKA, calcineurin and protein phosphatase 1 (PP1) [20,21]. The model of ref [9] for this subsection is based on the model proposed in Graupner & Brunel, 2007 [22]. We followed those models by assuming that postsynaptic plasticity is directly proportional to the calcium-dependent activation of CaMKII and set:

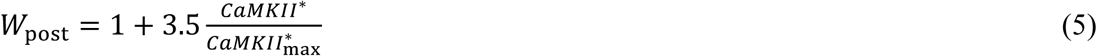

where *CaMKII*\* and 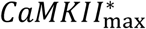 are the current concentration of activate (phosphorylated) CaMKII and its maximum value.

Our model also accounts for the biochemical pathways leading to the production of the endocannabinoids 2-arachidonoylglycerol (2-AG) and AEA, and their subsequent activation of cannabinoid receptors type-1, CB_1_R (see S1 Text for further details). 2-AG and AEA are retrograde signaling molecules that are produced in the postsynaptic neuron and diffuse to the presynaptic cell, where they activate CB_1_R [23,24,25]. We modeled CB_1_R activation by 2-AG and AEA using a simple three-state kinetic model (open, desensitized, inactivated), from which the open fraction *x*_CB1R_ was used to compute CB_1_R activation as

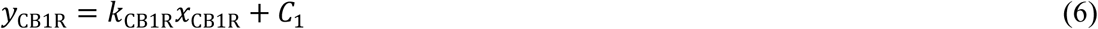

where *C*_1_ is a constant that accounts for the modulation of presynaptic plasticity by other pathways and *k*_CB1R_ quantifies the strength of CB_1_R activation on presynaptic plasticity. In our model, CB_1_R activation (*y*_CB1R_) controls the presynaptic weight *W*_pre_ according to the following rule: *W*_pre_ decreases for intermediate values of *y*_CB1R_, i.e. when *y*_CB1R_ is comprised between two tLTD thresholds 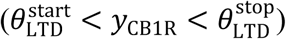 whereas *W*_pre_ increases when *y*_CB1R_ is larger than a tLTP threshold 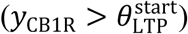. Following Cui et al., 2016, we implemented this rule as

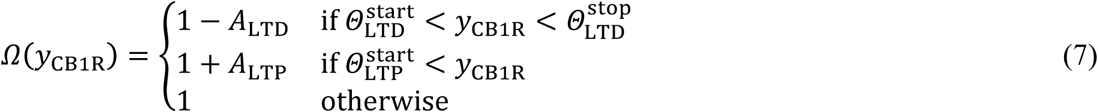

and

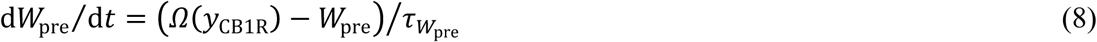

Here, *Ω* determines the change of presynaptic plasticity, with a time scale 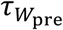 set to yield rapid changes of *W*_*pre*_ for large *y*_CB1R_values and very slow changes at very low *y*_CB1R_ (see S1 Text). To account for experimental observation that the presynaptic weight ranges from about 50 to 300%, *W*_*pre*_ was clipped to 3.0 (hard bound).

#### Model implementation and numerics

Our mathematical model comprises 36 ordinary differential equations and close to 150 parameters, among which more than one half is constrained by experimental data. Numerical solution was obtained with the LSODA solver from the ODEPACK fortran77 library with absolute and relative tolerances both equal to 10^−7^. Initial conditions were set to the steady-state of each variable in the absence of stimulation. Numerical integration proceeded until the synaptic weights reach stable values (typically observed around t ≈ 5min after the end of the stimulation protocol). We used the final values of the pre- and postsynaptic weights to compute the total synaptic weight change due to the stimulations. Importantly, with the exception of the stimulation protocols, *we have used the exact same equations and parameter values as in* [9]. The list of parameters and their estimated values is given in S2 Table. The current study employs stochastic simulations since the stimulation protocol is stochastic (while the rest of the model is deterministic). Therefore, the model was calibrated using experimental data from deterministic stimulation protocols [9] and we test here whether it can make successful predictions when we apply stochastic stimulation protocols, for which it was not calibrated. The results we present are thus averages over *N*_trial_ realizations. *N*_trials_ was varied from 30 to 500 depending on the smoothness of the averaged curves and values of SEM, but *N*_trials_=50 in most of the simulations. The computer code of the model is available online from https://github.com/iprokin/Cx-Str-STDP and https://senselab.med.yale.edu/modeldb/ShowModel.cshtml?model=187605.

## RESULTS

Corticostriatal synapses exhibit a bidirectional STDP in which NMDAR-tLTP, eCB-tLTD [15,16,17,18,19,26] or eCB-tLTP [8,9] are induced depending on the number of pairings (*N*_pairings_) and the spike timing (Δ*t*_STDP_). While STDP is classically investigated using fixed Δ*t*_STDP_, we examine here the effect of noisy Δ*t*_STDP_, which is more likely to happen in *in vivo*-like firing. We note Δ*t*_STDP_<0 when the postsynaptic stimulation occurred before the paired presynaptic one (post-pre pairings), whereas Δ*t*_STDP_ >0 when the presynaptic stimulation occurred before the postsynaptic one (pre-post pairings) (Fig. 1A).

**Figure 1:**
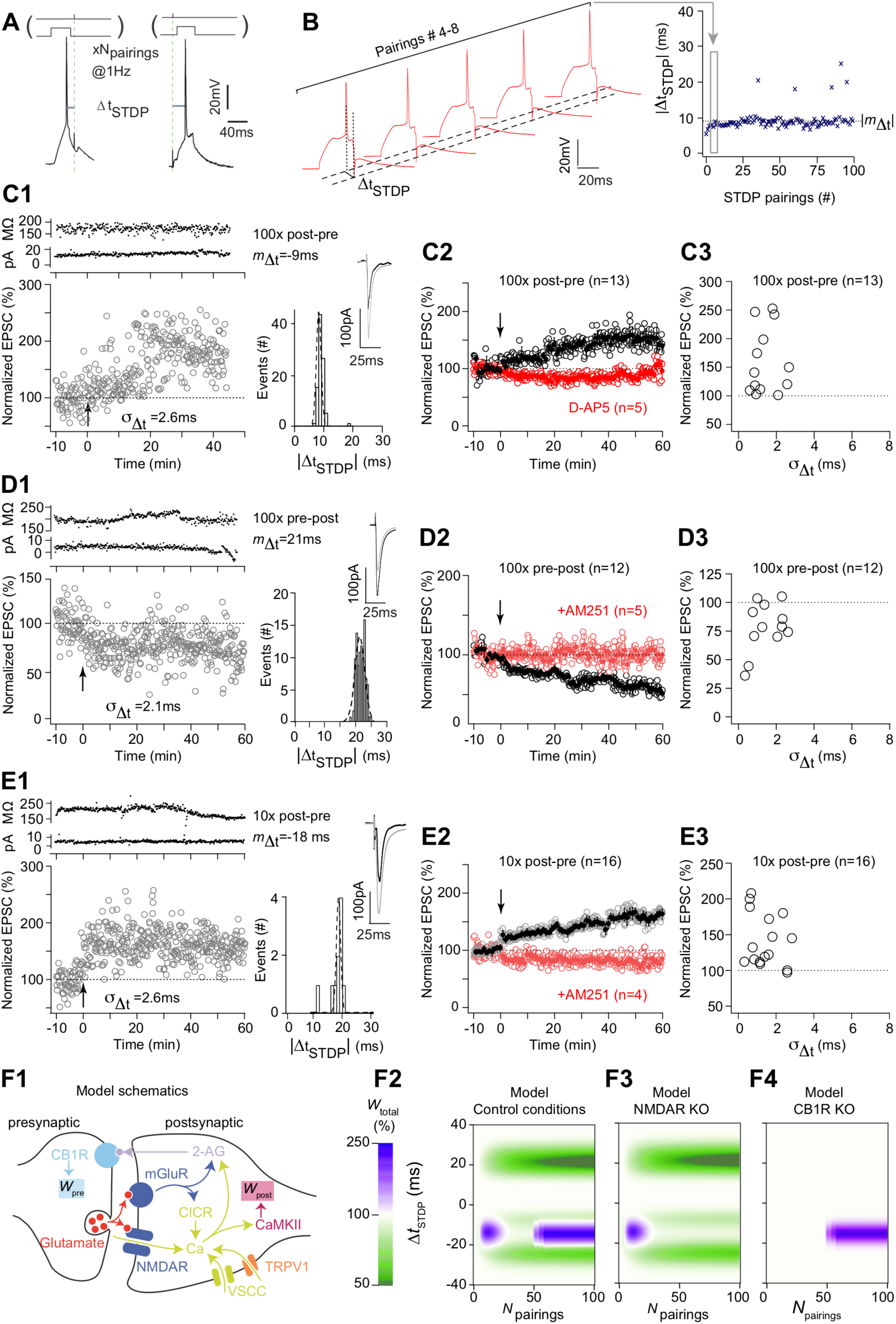
NMDAR- and eCB-mediated STDP. (**a**) STDP protocol: a single spike evoked by a depolarizing current step in the recorded striatal MSN was paired with a single cortical stimulation; this pairing was repeated 10 or 100 (*N*_pairings_) times at 1 Hz. Δ*t*_STDP_ indicates the time between pre- and postsynaptic stimulations. Δ*t*_STDP_ <0 and Δ*t*_STDP_ >0 refer to post-pre and pre-post pairings, respectively. (**b**) Example of 5 successive pairings (#4–8) (taken from the experiment illustrated in C1) exhibiting relatively fixed values of Δ*t*_STDP_ (#4: −8.0, #5: −7.8, #6: −7.4, #7: −8.8 and #8: −8.0 ms); these pairings are indicated with the grey frame within the 100 pairings. Plotting the successive Δ*t*_STDP_ illustrates the low value of their variance σ_Δ*t*_^2^ (2.6 ms). (**c**) Corticostriatal NMDAR-mediated t-LTP. (**c1**) Example of tLTP induced by 100 post-pre pairings. The statistics of Δ*t*_STDP_ were *m*_Δ*t*_ =-9 ms (mean) and σ_Δ*t*_ =2.6 ms (s.d.). Upper panel, time course of Ri (baseline: 172±2 MΩ and 50–60 min after pairings: 170±3 MΩ; change of 1%) and holding current (I_hold_) for this cell. Inset: distribution of Δ*t*_STDP_ for each of the 100 pairings. (**c2**) Averaged time-courses of tLTP induced by 100 post-pre pairings; this tLTP was mediated by NMDAR, because tLTP was prevented by the application of D-AP5 (50 µM). (**c3**) Relationship between the STDP magnitude and the STDP jitter for each of the recorded neurons (n=13). (**d**) Cortico-striatal eCB-mediated t-LTD. (**d1**) Example of tLTD induced by 100 pre-post pairings (*m*_Δ*t*_ =+21 ms with σ_Δ*t*_ =2.1 ms). Upper panel, time course of Ri (baseline: 195±1 MΩ and 50–60 min after pairings: 196±2 MΩ; change <1%) and I_hold_ for this cell. Inset: distribution of Δ*t*_STDP_ for the 100 pairings. (**d2**) Averaged time-courses of tLTD induced by 100 pre-post pairings; this tLTD was dependent on CB_1_R activation, because AM251 (3 µM) prevented tLTD. (**d3**) Relationship between the STDP magnitude and the jitter for each of the recorded neurons (n=12). (**e**) Cortico-striatal eCB-mediated tLTP. (**e1**) Example of tLTP induced by 10 post-pre pairings (*m*_Δ*t*_ =-18 ms with σ_Δ*t*_ =2.8 ms). Upper panel, time course of Ri (baseline: 205±1 MΩ and 50–60 min after pairings: 178±2 MΩ; change of 13%) and I_hold_ for this cell. Inset: distribution of Δ*t*_STDP_ for each of the 10 pairings. (**e2**) Averaged time-courses of tLTP induced by 10 post-pre pairings; this tLTP was dependent on CB_1_R activation, because AM251 prevented tLTD. (**e3**) Relationship between the STDP magnitude and the jitter for each of the recorded neurons (n=16). (**c1-d1-e1**) Insets correspond to the average EPSC amplitude at baseline (black trace) and at 40–50 min after STDP pairings (grey trace). (**c2-e2**) Error bars represent the SEM. (**f**) Predictions of the mathematical model in the absence of added jitter. The model expresses the biochemical pathways schematized in (**f1**): presynaptically released glutamate activates mGluR and NMDAR receptors. The latter gives rise to a transient of free cytosolic Ca, whereas the former generates endocannabinoid 2-AG and triggers calcium-induced calcium release (CICR), thus contributing to the Ca transient, together with VSCC and TRPV1. Free cytosolic calcium in turns activates the CaMKII/PKA/Calcineurin system and boosts 2-AG generation, thus increasing the activation of CB_1_R. In the model, the presynaptic weight is set directly by CB_1_R activation, whereas the postsynaptic weight is proportional to the fraction of activated CaMKII. See *Methods* and S1 Text for details. (**f2**) Changes of the total synaptic weight in the model in control conditions after paired stimulations with various Δ*t*_STDP_ and *N*_pairings_. The change in total synaptic weight, computed as the product of the pre- and postsynaptic weights, is given by the colorbar (green for tLTD, purple for tLTP). (**f3**) Removing the NMDAR/CaMKII/PKA/Calcineurin signaling from the model suppresses the tLTP observed for Δ*t*_STDP_ >0 and *N*_pairings_ > 50 in control conditions. The model thus reproduces NMDAR-tLTP. (**f4**) Conversely, removing CB_1_R from the model suppresses both the tLTP observed for Δ*t*_STDP_ >0 and *N*_pairings_ > 20 and the tLTD observed for Δ*t*_STDP_ <0. The model thus accounts for the experimental observations of eCB-tLTP and eCB-tLTD.

### NMDAR- and eCB-mediated STDP

Using the canonical paradigm for STDP, i.e. 100 pairings at 1 Hz we observed bidirectional STDP in MSNs for post- and presynaptic activities paired within −30< Δ*t*_STDP_ <+30 ms: 100 post-pre pairings induced tLTP whereas 100 pre-post pairings induced tLTD. An example of the tLTP induced by 100 post-pre pairings with fixed Δ*t*_STDP_ (Fig. 1B) is illustrated in Figure 1C1; the mean baseline EPSC amplitude was 95±3 pA before pairings, and increased by 77% to 168±6 pA 45 minutes after pairings. The input resistance Ri remained stable over this period. During a STDP experiment, the value of the successive Δ*t*_STDP_ slightly varies from one pairing to another (Fig.1B). This low jitter can be written formally as Δ*t*_STDP_ = *m*_Δ*t*_ + *ξ*_Δ*t*_, were *m*_Δ*t*_ is the mean spike timing and *ξ*_Δ*t*_ is a random variable with mean 0 and standard deviation *σ*_Δ*t*_. Note that in most experimental studies of STDP, Δ*t*_STDP_ and *m*_Δ*t*_ are not distinguished because *σ*_Δ*t*_ is expected to be low. In the STDP example of Figures 1B and 1C1, *m*_Δ*t*_ =-9 ms and the jitter is *σ*_Δ*t*_ =2.6 ms.

Overall, 100 post-pre pairings induced tLTP (mean EPSC amplitude recorded 60 min after protocol induction: 149±13% of baseline, *p*=0.0028, *n*=13; 11 of 13 cells displayed tLTP) (Fig. 1C2). *σ*_Δ*t*_ was in average 1.4±0.2 ms (Fig. 1C3). This tLTP was NMDAR-mediated since prevented with D-AP5 (50 µM), a NMDAR antagonist, (88±12%, *p*=0.375, n=5; 1/5 showed tLTP) (Fig. 1C2). Conversely, pre-post pairings induced tLTD, as shown in the example in Figure 1D1: the mean baseline EPSC amplitude, 206±4 pA, had decreased by 25%, to 154±8 pA, 45 minutes after pairings. Overall, 100 pre-post pairings induced tLTD (78±6%, *p*=0.0047, *n*=12; 9/12 cells displayed tLTD) (Fig. 1D2). *σ*_Δ*t*_ was in average 1.5±0.2 ms (Fig. 1D3). This tLTD was CB_1_R-mediated since prevented with AM251 (3µM), a CB_1_R specific inhibitor (99±2%, *p*=0.816, n=5; 0/5 showed tLTD) (Fig. 1D2). Using the canonical paradigm for STDP, we thus find an anti-Hebbian polarity for corticostriatal STDP. We previously showed that GABA operates as a Hebbian/anti-Hebbian switch at corticostriatal synapses [18,27] because corticostriatal STDP polarity depends on the presence of GABA_A_ receptor antagonists (*in vitro* Hebbian STDP, [15,16]) or not (*in vitro* anti-Hebbian STDP, [8,17,19]; *in vivo* anti-Hebbian STDP [26]). We thus recorded STDP in the absence of GABA_A_R antagonist to preserve the anti-Hebbian polarity as observed *in vivo* [26].

Besides this bidirectional STDP (NMDAR-tLTP and eCB-tLTD) induced for 100 pairings, we recently reported that low numbers of pairings (*N*_pairings_ =5-15) induce an eCB-tLTP, dependent on the activation of CB_1_R [8,9]. Figure 1E1 shows an example of tLTP induced by 10 post-pre pairings where the mean baseline EPSC amplitude was 159±3pA before pairings and has increased by 54% to 245±5pA 45 minutes after pairings. Overall, 10 post-pre STDP pairings induced tLTP (143±9%, p=0.0005, n=16; 14 cells out of 16 resulted in tLTP), which was prevented with AM251 (3µM) (76±10%, p=0.080, n=4; 0/4 showed tLTP) (Fig. 1E2) as recently reported [8,9]. *σ*_Δ*t*_ for 10 post-pre parings STDP was 1.7±0.2 ms (n=16) (Fig. 1E3). Note that there was no significant difference between *σ*_Δ*t*_ measured for NMDAR-tLTP, eCB-tLTD and eCB-tLTP (one way Anova, p=0.570).

To help interpret our experimental results, we used a mathematical model of the signaling pathways implicated in corticostriatal STDP, including NMDAR-dependent and CB_1_R-dependent plasticity [8,15,16,17]. The model is described in details in S1 Text, with parameter values listed in S2 Table. We also include in the Supporting Information a thorough description of the mechanisms by which eCB-dependent and NDMAR-dependent plasticities are expressed in the model when a canonical STDP protocol (*N*_pairings_=100) is applied (S3 Text). Figure 1F2 shows the value of the total synaptic weight *W*_total_ predicted by the model with various *N*_pairings_ and Δ*t*_STDP_ in control conditions. In agreement with the experimental data, with small *σ*_Δ*t*_ (3 ms), the model features three main plasticity regions. Short pre-post pairings (Δ*t*_STDP_>0) give rise to tLTD with smoothly increasing amplitude when *N*_pairings_ increases. For short post-pre pairings (Δ*t*_STDP_<0), the model exhibits two plasticity regions with similar amplitude: a first tLTP region is observed for low numbers of pairings (3<*N*_pairings_<25) while a second tLTP region is expressed only when *N*_pairings_>50. Blocking the NMDAR-CaMKII pathway in the model (Fig. 1F3), suppresses the second tLTP region (Δ*t*_STDP_ <0; *N*_pairings_>50) whereas blocking CB_1_R activation (Fig. 1F4) prevents the expression of tLTD (Δ*t*_STDP_ >0; *N*_pairings_>15) and of the first tLTP (Δ*t*_STDP_<0; 3<*N*_pairings_<25). Therefore, the model emulates both CB_1_R-dependent tLTD and tLTP, in agreement with the eCB-tLTD and eCB-tLTP illustrated in the experimental data of Figure 1D and 1E, respectively. The model also faithfully emulates the appearance of NMDAR-tLTP for *N*_pairings_ >50.

### Predicting the effects of jittered STDP with a mathematical model

The small jitter observed in the above experiments was the result of biological variety inherent to the experimental setup. Our next goal was to extend this study to larger jitters (3<*σ*_Δ*t*_<10 ms) that we controlled experimentally. We first preserved determinism for the interval between two consecutive pairings (Inter-Pairing Interval, IPI) and used the same definition as above for jittered Δ*t*_STDP_ (see *Methods*): Δ*t*_STDP_ = *m*_Δ*t*_ + *ξ*_Δ*t*_, where *ξ*_Δ*t*_ is a random variable with zero mean and *m*_Δ*t*_ is the expected value of the spike timing Δ*t*_STDP_ over the *N*_pairings_. In the following, we adjusted the probability distribution function and the variance of *ξ*_Δ*t*_ to perform a parametric exploration of the impact of jitter.

In Figure 2, the jitter *ξ*_Δ*t*_ was sampled from a uniform distribution in [-*σ*_Δ*t*_,+ *σ*_Δ*t*_] (Fig. 2A1). With *σ*_Δ*t*_ = 0, i.e. with no added noise, the model reproduces the same results as Figure 1C above, exhibiting eCB-tLTD for *m*_Δ*t*_ > 0 and large enough *N*_pairings_, and eCB-tLTP / NMDAR-tLTP for *m*_Δ*t*_ < 0 and low / large values of *N*_pairings_, respectively (Fig. 2B1).

**Figure 2:**
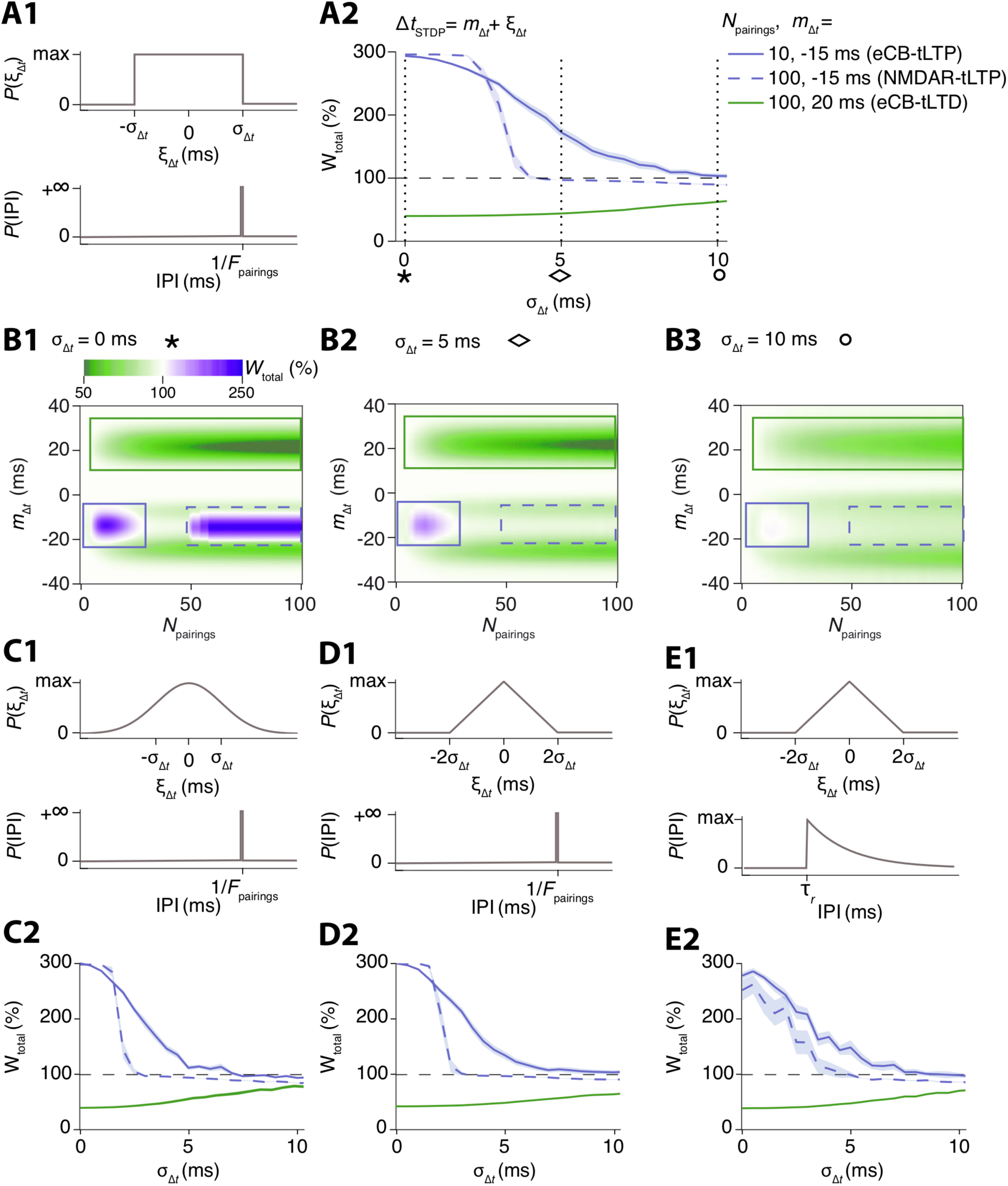
The mathematical model predicts that NMDAR-tLTP is less robust to jitter than eCB-tLTP and eCB-tLTD. (**a**) Using a jittered STDP protocol combining uniformly-distributed jitter of the spike timing with deterministic IPIs at 1 Hz (**a1**), the amplitudes of the three plasticities of the model decrease with increasing jitter amplitudes *a*_*At*_ (**a2**). eCB-tLTP (purple full line) was triggered with 10 pairings of average spike timing *m*_*At*_ = −15 ms and eCB-tLTD (green full line) with 100 pairings of *m*_*At*_ = +20 ms. NMDAR-tLTP (purple dashed line) corresponded to 100 pairings of *m*_*At*_ = −15 ms. eCB-tLTD is weakly affected by the jitter whereas eCB-tLTP degrades at a faster rate. NMDAR-tLTP is less robust. Full and dashed lines show averages and light swaths show ±1 sem. (**b**) Evolution of the three plasticity domains foru_At_ =0 (**b1**), 5 (**b2**) and 10 ms (**b3**) for the STDP protocol of (**a**). Note the persistence of the eCB-tLTD domain (green box) while eCB-tLTP (purple box) and NMDAR-tLTP domain (purple dashed box) disappeared for 5 and 10 ms, respectively. (**c-e**) Similar results are obtained with other jitter distributions, including Gaussian-distributed jitter (**c**) and triangularly-distributed jitter (**d**). We also considered a combination of triangularly-distributed jitter with random IPIs distributed according to an exponential distribution with refractory period T_r_=0.95 s (see *Methods*) in (**e**). The plots of the amplitude of the 3 plasticities for increasing jitter amplitude *a*_*At*_ (**c2, d2, e2**) show similar profiles and are similar to panel (**a2**), indicating that the model predictions are not qualitatively modified by the probability distribution function of the jitter.

STDP in the model was globally not robust to jitter, since the three forms of plasticity vanished when *σ*_Δ*t*_ was large enough (Fig. 2A2). However, the model delivered the prediction that the three forms of plasticity are not equally susceptible to jitter. In particular, NMDAR-tLTP is predicted to be very fragile when subjected to *σ*_Δ*t*_: its amplitude decreases as soon as *σ*_Δ*t*_ > 2 ms and completely vanishes for *σ*_Δ*t*_ > 4 ms (Fig. 2A2 and 2B2). In contrast, STDP-tLTP is much less sensitive: eCB-tLTP is still present for jitters as large as 7–8 ms and eCB-tLTD is still expressed with *σ*_Δ*t*_ = 10 ms (Fig. 2B3).

We then evaluated the validity of this prediction when the probability distribution function of the jitter changes. Figure 2 illustrates the results we obtained with deterministic IPI but Gaussian (Fig. 2C) or triangular (Fig. 2D) distributions of *σ*_Δ*t*_ (see *Methods*). In both cases, the profiles of the robustness curves (Fig. 2C2 and 2D2) are almost identical to each other and similar to the robustness curves obtained with an uniform distribution (Fig. 2A2). In Figure 2E, we combined a triangular distribution for *σ*_Δ*t*_ with random IPIs, i.e. in this case, stochasticity is applied not only to the timing between the two stimulations of a given pairing, but also to the time interval between two consecutive pairings: IPI distribution was Poisson with refractory period *τ*_*r*_ and rate *λ* (see *Methods*). Despite this increased stochasticity, we found that STDP-tLTD is again more robust, tolerating jitters up to 6-7 ms (eCB-tLTP) or 10 ms (eCB-tLTD) whereas NMDAR-tLTP is fragile, vanishing for σ_Δ*t*_ > 4 ms (Fig. 2E2). Therefore, according to our model, the observation that eCB-STDP is more robust to jitter than NMDAR-tLTP should be a generic property of the response of the signaling pathways (NMDAR- *versus* eCB-mediated plasticity) to noisy paired stimulations and should not be crucially dependent on the probability distribution function of the noise.

### Differential effect of jitter onto NMDAR- and eCB-mediated plasticity

Based on the model predictions, we next investigated experimentally the sensitivity to jitter of NMDAR-tLTP induced by 100 post-pre pairings (Fig. 3A), using the distribution of Δ*t*_STDP_ and IPI shown in the Figure 2C1. We set the range of σ_Δ*t*_ values explored to [0,10] ms because the model predicts that this range is enough to obliterate the three plasticities (Fig. 2A2). As aforementioned, NMDAR-tLTP can be induced with a σ_Δ*t*_<3 ms (Fig. 1C3). Increasing σ_Δ*t*_ to 3-10 ms was sufficient to obliterate tLTP for 100 post-pre pairings. As exemplified (Fig. 3B and C1), 100 post-pre pairings with σ_Δ*t*_ = 7.9 ms and centered on *m*_Δ*t*_ = −18 ms failed to induce significant plasticity (the mean baseline EPSC amplitude, 194±3 pA, did not show significant change one hour after pairing, 186±4 pA). Overall, we did not observe tLTP expression for σ_Δ*t*_ > 3 ms (92±2%, *p*=0.560, n=7; 0/7 showed tLTP) (Fig. 3C2 and C3) in agreement with the model prediction (Fig. 2C2). Therefore, with deterministic IPI=1sec, NMDAR-tLTP appeared indeed fragile to jittering.

**Figure 3:**
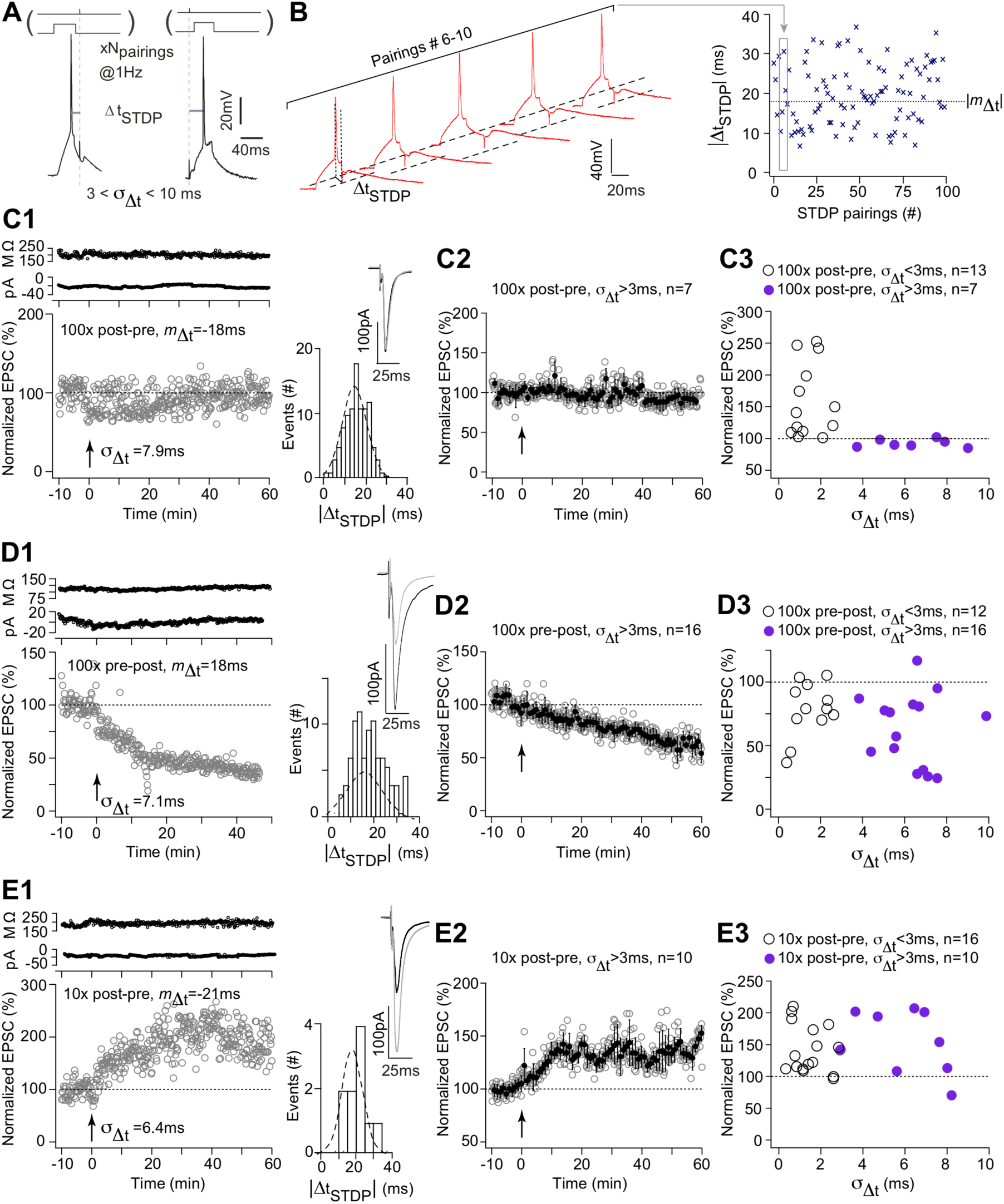
Differential effect of jitter onto NMDAR- and eCB-mediated plasticity. (**a**) STDP protocol; Δ*t*_STDP_ indicates the time between pre- and postsynaptic stimulations. Δ*t*_STDP_ <0 and Δ*t*_STDP_ >0 refer to post-pre and pre-post pairings, respectively. (**b**) Example of 5 successive pairings (#5–9) (taken from the experiment illustrated in c1) exhibiting jittered values of Δ*t*_STDP_ (#5: −10.3, #6: −15.2, #7: −30.5, #8: −20.7 and #9: −17.3 ms); these pairings are indicated with the grey frame within the 100 pairings displaying jittered Δ*t*_STDP_. The distribution of the successive Δ*t*_STDP_ illustrate their standard deviation *a_At_=7.9* ms. (**c**) NMDAR-mediated t-LTP is prevented with σ_Δ*t*_>3 ms. (**c1**) Example of the absence of plasticity after 100 post-pre pairings with *m*_Δ*t*_ =-18 ms and *a_At_=7.9* ms. Upper panel, time course of Ri (baseline: 198±2 MΩ and 50–60 min after pairings: 194±2 MΩ; change of 2%) and holding current (Iho_ld_) for this cell. Inset: distribution of Δ*t*_STDP_ for the 100 pairings. (**c2**) Averaged time-courses of the absence of plasticity after 100 post-pre pairings with 3<σ_Δ*t*_<10 ms. (**c3**) Relationship between the STDP magnitude and the jitter for each of the recorded neurons (n=20). (**d**) eCB-mediated t-LTD is not affected by 3<σ_Δ*t*_<10 ms. (**d1**) Example of tLTD induced by 100 pre-post pairings with m_At_=+18 ms and *a_At_=7.1* ms. Upper panel, time course of Ri (baseline: 112±1 MΩ and 50–60 min after pairings: 118±1 MΩ; change of 5%) and Iho_ld_ for this cell. Inset: distribution of Δ*t*_STDP_ for the 100 pairings. (**d2**) Averaged time-courses of tLTD induced by 100 pre-post pairings with 3<σ_Δ*t*_<10 ms. (**d3**) Relationship between the STDP magnitude and the jitter for each of the recorded neurons (n=28). (**e**) eCB-mediated tLTP is not affected by 3<σ_Δ*t*_<10 ms. (**e1**) Example of tLTP induced by 10 post-pre pairings with m_At_=-21 ms and *a_At_=6*.4 ms. Upper panel, time course of Ri (baseline: 185±1 MΩ and 50–60 min after pairings: 197±2 MΩ; change of 6%) and I_hold_ for this cell. Inset: distribution of Δ*t*_STDP_ for the 10 pairings. (**e2**) Averaged time-courses of tLTP induced by 10 post-pre pairings with 3<σ_Δ*t*_<10 ms. (**e3**) Relationship between the STDP magnitude and the jitter for each of the recorded neurons (n=26).Insets correspond to the average EPSC amplitude at baseline (black trace) and at 40–50 min after STDP pairings (grey trace). Error bars represent the SEM.

We then similarly investigated the robustness of the eCB-tLTD to the jitter of Δ*t*_STDP_. We observed potent tLTD even with large values of σ_Δ*t*_. Indeed, as shown in Figure 3D1, 100 pre-post pairings with σ_Δ*t*_ = 7.0 ms centered on *m*_Δ*t*=_ +18 ms induced a large tLTD: the mean baseline EPSC amplitude, 160±3 pA, had decreased by 50%, to 81±5 pA, 45 minutes after pairings. Overall, we observed that tLTD was still induced for 100 pre-post pairings with 3 <σ_Δ*t*_< 10 ms (70±7%, *p*=0.002, n=16; 15/16 showed tLTD) (Fig. 3D2 and D3). The mean value of tLTD, observed for 3 <σ_Δ*t*_< 10 ms, was not significantly different from the one observed in control, i.e. for 0 <σ_Δ*t*_< 3 ms (*p*=0.400). Therefore, as predicted by our mathematical model, the experiments show that eCB-tLTD appears very robust to the jitter of Δ*t*_STDP_, since eCB-tLTD resists to the largest σ_Δ*t*_ values used in this study.

Finally, we investigated the robustness of the eCB-tLTP to the jitter of Δ*t*_STDP_ and observed potent tLTP even with large values of jitter, i.e. up to σ_Δ*t*_ ~ 8 ms. An example of tLTP induced by 10 post-pre pairings with σ_Δ*t*_ = 6.4 ms centered on *m*_Δ*t*_ = −21 ms is illustrated in Figure 3E1: the mean baseline EPSC amplitude was 113±2 pA before pairings, and increased by 115% to 243±6 pA 45 minutes after pairings. In summary, tLTP could be induced even for σ_Δ*t*_ = 8 ms (165±9%, p=0.0007, n=10; 8/10 showed tLTP) (Fig. 3E2 and E3); the mean value of tLTP was not significantly differently than the one observed in control (*p*=0.166). Therefore, eCB-tLTP appears to be robust to the jitter of Δ*t*_STDP_ for σ_Δ*t*_ up to 8 ms, in quantitative agreement with our model predictions (Fig. 2).

We ensured that the *m*_Δ*t*_ values were not significantly different in control and jittered conditions for 100 post-pre pairings (18±2 ms, n=13, *vs* 21±3 ms, n=7, p=0.4711), 100 pre-post pairings (21±2 ms, n=12, *vs* 20±2 ms, n=16, p=0.6204), 10 post-pre pairings (16±2 ms, n=16, *vs* 21±1 ms, n=10, p=0.0788) or between these different STDP protocols (oneway ANOVA: p=0.1140) (S4 Figure).

In conclusion, whereas NMDAR-mediated tLTP is very fragile against the jitter of Δ*t*_STDP_, eCB-mediated plasticity (eCB-tLTD as well as eCB-tLTP to a lesser degree) exhibits a large robustness for the temporal imprecision of Δ*t*_STDP_.

### Model-based analysis of the effects of jittering

The above experimental results provide us with a validation of our mathematical model. We emphasize that the values of the model parameters were estimated using experimental data obtained with regular (deterministic) stimulation protocols [9] and we used the model here with stochastic protocols, without any adaptation or change of parameter values. Hence, although the model was calibrated with deterministic stimulations, it provides successful predictions with stochastic stimulations for which it was not calibrated. We next used this validated model as a tool to investigate the molecular mechanisms behind these differences of robustness.

We have shown in a previous study that eCB-tLTP is expressed when the amount of eCBs produced is large enough that the fraction of activated CB_1_R, *y*_CB1R_, overcomes a threshold 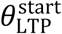 [9]. In the absence of added noise, σ_Δ*t*_ = 0, our experimental data shows that as few as 5 pairings at 1Hz and Δ*t*_STDP_ = −15 ms are enough to trigger eCB-tLTP [8]. Accordingly, our model produces large amounts of eCBs in the first 5-10 pre-post pairings so that *y*_CB1R_ reaches 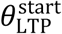 after as few as 5 pairings (Fig. 4A1, purple). When σ_Δ*t*_ =5 ms jitter is added to *m*_Δ*t*_ = −15 ms, some of the pairings fail to deliver *y*_CB1R_ transients of maximal amplitude (Fig. 4A2). But for σ_Δ*t*_< 7-8 ms, a number of *y*_CB1R_ transients still have sufficient amplitude to overcome 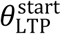 which is enough to trigger eCB-tLTP (Fig. 4A2). When σ_Δ*t*_ is large, though (e.g. 10 ms), the *y*_CB1R_ transients fail to reach the LTP zone and eCB-tLTP is not expressed (Fig. 4A3).

In experiments without added jitter, eCB-tLTD progressively accumulates when *N*_pairings_ increases and starts to be significant for *N*_pairings_ >25 [8]. In the model with *m*_Δ*t*_ = +20 ms and σ_Δ*t*_ =0, the *y*_CB1R_ transients remain in the LTD area for the most part of the STDP pairings (Fig. 4A3 light green). To account for the progressive accumulation of eCB-tLTP with *N*_pairings_, each transient in the model contributes a small decrease of the synaptic weight. Even with large amounts of jitter (see e.g. σ_Δ*t*_ =5 or 10 ms in Figure 4A2-3) *y*_CB1R_ transients remain mostly inside LTD area so eCB-tLTD remains expressed even with large jitter.

**Figure 4:**
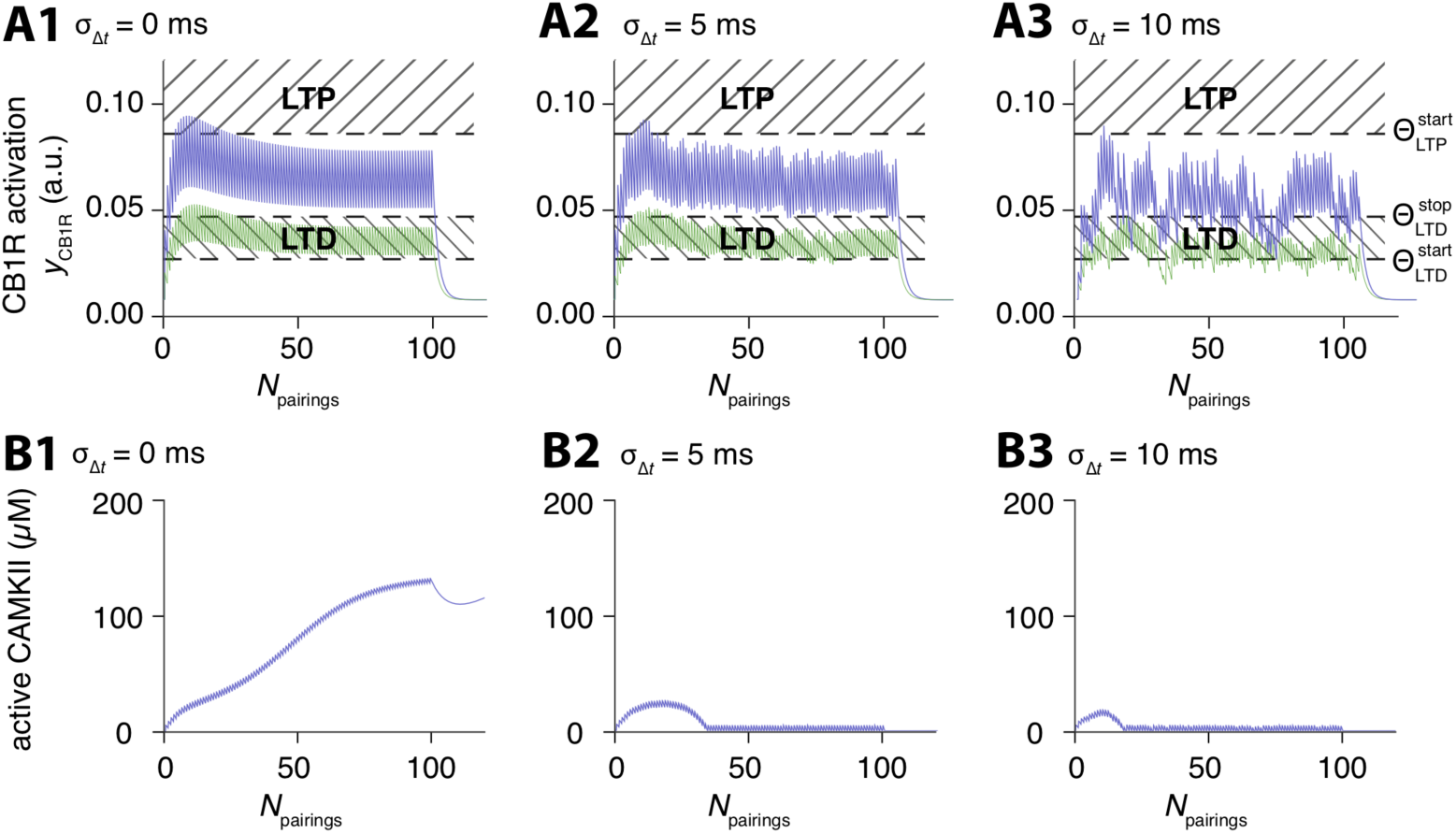
The mathematical model provides an explanation for the differential robustness of plasticity. Model-predicted time-courses of (**a**) CB1R activation *y*CB1R and (**b**) the concentration of active CaMKII during a protocol with jitter amplitude σ_Δ*t*_ = 0 (**a1-b1**), 5 (**a2-b2**) or 10 (**a3-b3**) ms. The STDP protocol was the same as Figure 2a1 i.e. uniformly-distributed jitter of the spike timing and deterministic IPIs at 1 Hz. eCB-tLTP (**a**, purple full line) and NMDAR-tLTP (**b**, purple dashed line) were triggered with average spike timing *m*_Δ*t*_ = −15 ms and eCB-tLTD (**a**, green full line) with *m*_Δ*t*_ = +20 ms. In (**a**) the shaded boxes locate the areas where *y*CB1R triggers eCB-tLTP or eCB-tLTD. Those areas are defined by the thresholds 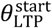 and 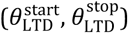, respectively (see *Materials and Methods*). For *m*_Δ*t*_ = −15 ms (purple), increasing jitter amplitudes progressively hinders the build up of the *y*CB1R transients. *y*CB1R still reaches the LTP area for 5–25 pairings with σ_Δ*t*_ = 0 (**a1**) and 5 ms (**a2**) thus triggering eCB-tLTP but fails to do so for σ_Δ*t*_ = 10 ms (**a3**). Conversely, eCB-tLTD is robust to jitter because *y*CB1R remains in the LTD region for most of the pairings with *m*_Δ*t*_ = +20 ms (green), for all tested jitter amplitudes. In (**b**), with *m*_Δ*t*_ = −15 ms (purple) the switch from the low activation to the high activation state of CaMKII is obtained in the absence of jittering σ_Δ*t*_ = 0 ms (**b1**). The progressive build up of activated CaMKII is very efficiently suppressed as soon as σ_Δ*t*_ = 5 ms (**b1**) thus effectively suppressing NMDAR-tLTP.

### The fragility of NMDAR-tLTP is predicted to be frequency-dependent

Experimentally, NMDAR-tLTP (*m*_Δ*t*_ = −15 ms and σ_Δ*t*_ =0) is observed at *F*_pairings_ = 1 Hz when *N*_pairings_ >50, beyond which its amplitude does not depend much on *N*_pairings_ [9]. To account for this feature, the steady-state concentration of activated CaMKII, that sets *W*_post_ in the model, is bistable: 45 min after the stimulation, CaMKII is either almost fully inactivated (“no plasticity” state) or almost fully activated (“LTP” state) [9,22]. When the frequency of post-pre pairings (−30< Δ*t*_STDP_<0 ms) is large enough (i.e. *F*_pairings_ ≥ 1 Hz), the IPI is smaller than the decay time of the CaMKII activation transient triggered by each post-pre pairings (Fig. 4B1). As a result, in the absence of added noise (σ_Δ*t*_ =0), the CaMKII activation transients progressively build up on top of each other. The accumulated CaMKII activation overcomes the threshold between the “no plasticity” and the “LTP” states only for *N*_pairings_ >50 (Fig. 4B1, purple) (see [9]). With jitter (e.g. σ_Δ*t*_ =5 ms in Fig. 4B2 or 10 ms in Fig. 4B3), many of the IPIs are either too long or too short to trigger maximal amplitude transients of activated CaMKII. As a result, CaMKII activation never reaches the threshold. The trajectories remain in the basin of attraction of the “no plasticity” state and tLTP cannot be expressed, thus explaining the fragility of NMDAR-tLTP with respect to jittering (at 1 Hz).

A major indication from the above analysis is the importance of the IPI frequency for the robustness of NMDAR-tLTP to jitter. Indeed, with smaller IPI (/larger pairing frequencies), the stimulations would be more efficient to bring CaMKII activation transients above the LTP threshold compared to the 1 sec (/1Hz) (Fig. 4B1). Figure 5 compares the robustness to jitter for deterministic IPI with *F*_pairings_=1 Hz (Fig. 5A) and 1.05 Hz (Fig. 5B). The size of the green area that locates eCB-tLTD on Figure 5A2-B2 is not much altered by the increase of the pairing frequency. Likewise, the size of the purple area that locates eCB-tLTP in Figure 5A1-B1 does not vary much when the frequency is increased. However, the size of the purple area locating NMDAR-tLTP increases drastically with frequency: whereas it stops around σ_Δ*t*_ = 4 ms for *F*_pairings_ =1Hz (Fig. 5A2), it extends beyond σ_Δ*t*_ =10 ms when *F*_pairings_ is increased (Fig. 5B2). Therefore, our model predicts that NMDAR-tLTP should become more robust to jitter at larger frequencies, whereas eCB-tLTP and eCB-tLTD are not significantly changed.

**Figure 5:**
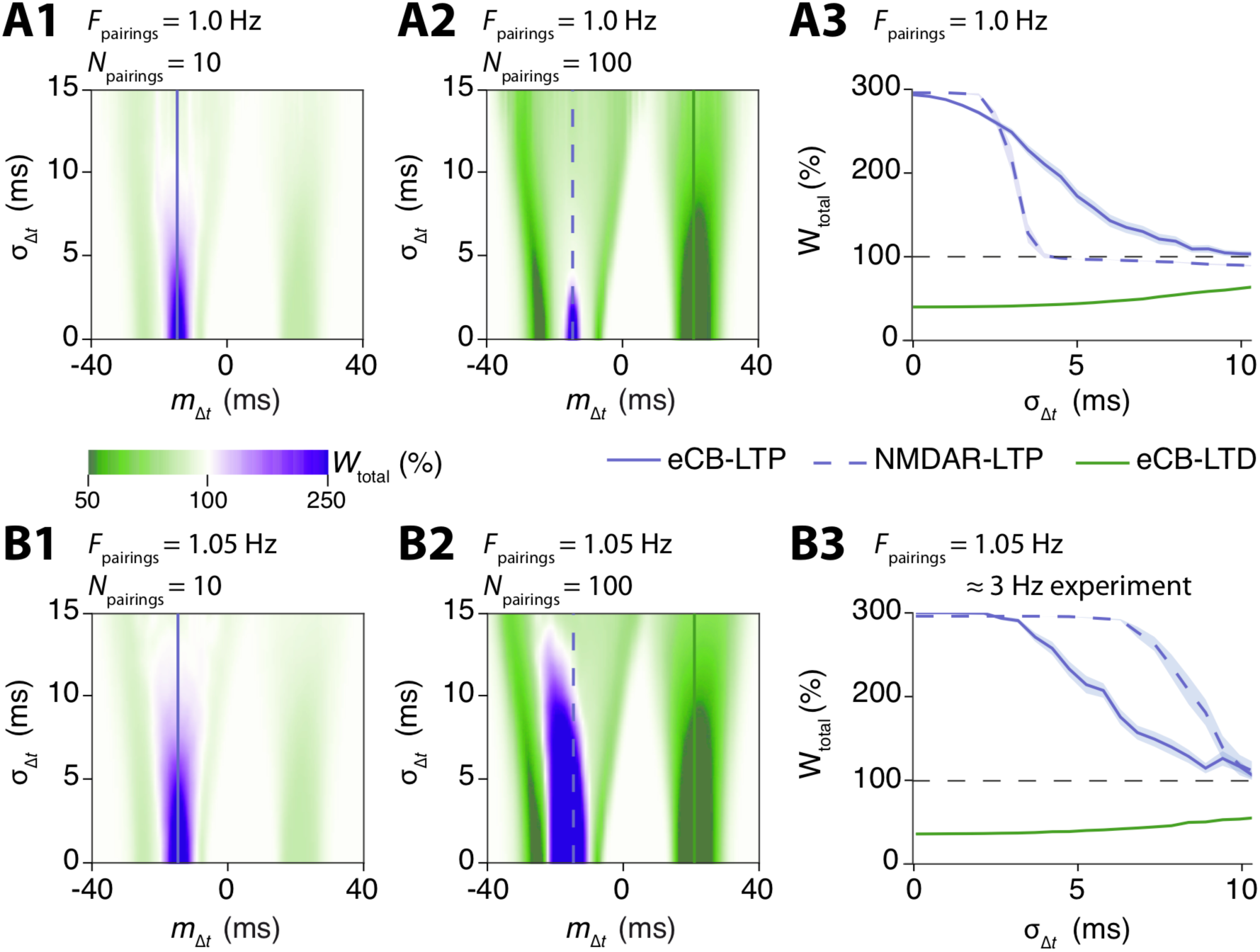
The model predicts a change of robustness pattern when frequency is increased. (**a**) Summary of the robustness of the model plasticity at 1 Hz. The jittered STDP protocol combined uniformly-distributed jitter of the spike timing and deterministic IPIs at 1 Hz. Two-dimensional maps show alterations of the plasticities obtained with 10 (**a1**) or 100 (**a2**) pairings, when the average spike timing *m*_Δ*t*_ and the jitter amplitude σ_Δ*t*_ are changed. The locations of the plasticity blotches are indicated with horizontal lines: full purple for eCB-tLTP, full green for eCB-tLTD and dashed purple for NMDAR-tLTP. The length of each of these blotches along the y-axis reflects their robustness to jitter. (**a3**) A cross-section along the dashed and full horizontal lines of (**a1-a2**) illustrates the robustness of the three plasticities. (**b**) Robustness of the model with larger frequency (1.05 Hz) and same jittered STDP protocol as in (**a**). Note that there exists a distortion between the pairing frequency in the model and in the experiments. Here, the behavior of the model for *F*_pairings_=1.05Hz corresponds to experimental results with *F*_pairings_ ≈3Hz **[9]**. The two-dimensional maps for 10 (**b1**) or 100 (**b2**) pairings as well as the cross-sections (**b3**) predicts that for frequencies larger than 1Hz, the robustness pattern changes, with NMDAR-tLTP becoming more robust.

Several aspects of glutamate signaling at the corticostriatal synapse are known to display complex frequency-dependence. Examples include the amount of glutamate released upon each presynaptic spike, glutamate uptake by transporters or glutamate receptor activation due to desensitization of AMPAR [28]. Our model is calibrated with experimental data at 1 Hz and features none of the above frequency dependencies. Therefore, we cannot expect a precise quantitative match between experiments and model when frequency is varied below or above 1Hz. In particular, sensitivity to frequency changes is much larger in the model than in the experiments [9]. However, the model still yields correct predictions of the main qualitative trends observed in the experiments, since the effects of a small change of *F*_pairings_ in the model (1.00 to 1.05 Hz) are similar to the effects of larger changes (1 to 3 Hz) in the experiments [9]. Therefore, in the experiments, we expect to observe a change of robustness pattern similar to Figure 5B, around *F*_pairings_=3 Hz rather than 1.05 Hz.

### Increasing *F*_pairings_ and *N*_pairings_ stabilized NMDAR-tLTP against jitter

We next tested experimentally whether an increase from 1 to 3 Hz of STDP pairings would protect the NMDAR-tLTP against jitter, as predicted by the model. In these conditions, we observed tLTP even with large jitters. Indeed, as shown in Figure 6A1, 100 post-pre pairings at 3 Hz with σ_Δ*t*_ = 9 ms and centered on *m*_Δ*t*_ =-22 ms induced tLTP: the mean baseline EPSC amplitude, 194±4 pA, had increased by 212%, to 606±7 pA, 45 min after pairings. Overall, we observed a significant tLTP for 100 post-pre pairings applied at 3 Hz with 4<σ_Δ*t*_<9 ms (173±3%, p=0.02, n=10; 6/10 showed tLTP) (Fig. 6A2 and A3); the mean value of these tLTP observed at 3 Hz with 4<σ_Δ*t*_<9 ms were not significantly different from the one observed with stimulation-related σ_Δ*t*_< 3 ms at 1 Hz (p=0.520). When we plotted the plasticity magnitude against σ_Δ*t*_ for 100 post-pre pairings at 3 Hz, we observed that tLTP was still induced even for σ_Δ*t*_ = 9 ms (Fig. 6A3). The NMDAR-mediated tLTP thus acquires certain robustness to the jitter of Δ*t*_STDP_ with increasing frequency of pairings.

**Figure 6:**
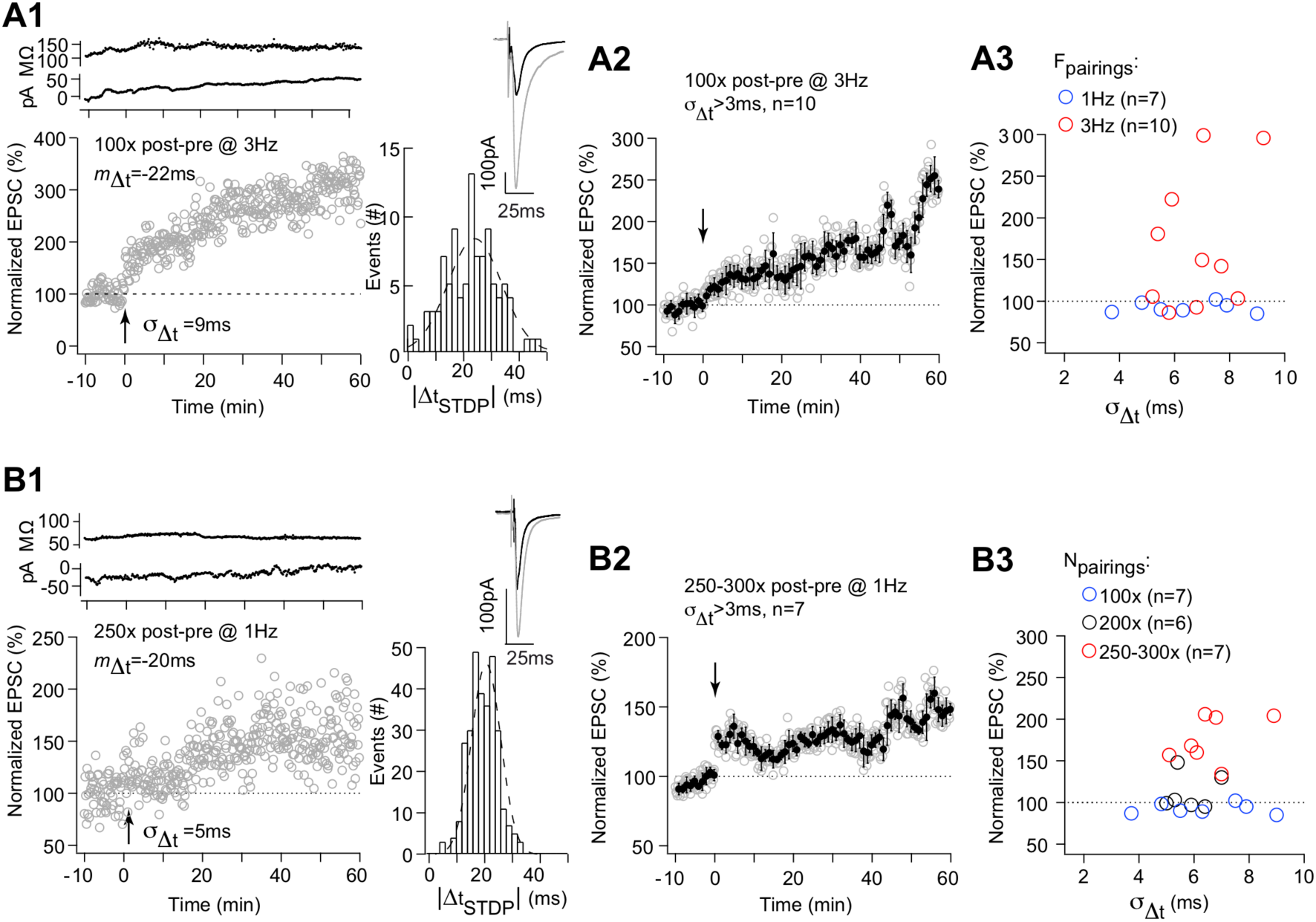
Increasing frequency and *N*_pairings_ stabilized NMDAR-tLTP against jitter. (**a**) 100 post-pre pairings at 3 Hz induced a tLTP which is not affected by 3<σ_Δ*t*_^2^ <10 ms. (**a1**) Example of tLTP induced by 100 post-pre pairings at 3 Hz with *m*_Δ*t*_ =-22 ms and σ_Δ*t*_^2^=9 ms. Bottom, time course of Ri (baseline: 123±1 MΩ and 50–60 min after pairings: 138±1 MΩ; change of 12%) and I_hold_ for this cell. Inset: distribution of Δ*t*_STDP_ for the 100 pairings. (**a2**) Averaged time-courses of tLTP induced by 100 post-pre pairings at 3 Hz with 3<σ_Δ*t*_^2^<10 ms. (**a3**) Relationship between the STDP magnitude and the jitter for each of the recorded neurons after 100 pairings at 1 Hz (n=7; blue circles) or at 3 Hz (n=10; red circles). (**b**) 250–300 post-pre pairings induced tLTP which is not affected by 3<σ_Δ*t*_^2^ <10 ms. (**b1**) Example of tLTP induced by 250 post-pre pairings with *m*_Δ*t*_ =-20 ms and σ_Δ*t*_^2^=5 ms. Bottom, time course of Ri (baseline: 65±1 MΩ and 50–60 min after pairings: 65±2 MΩ; change <1%) and Ihold for this cell. Inset: distribution of Δ*t*_STDP_ for the 250 pairings. (**b2**) Averaged time-courses of tLTP induced by 250–300 post-pre pairings with 3<σ_Δ*t*_^2^ <10 ms. (**b3**) Relationship between the STDP magnitude and the jitter for each of the recorded neurons after 100 (n=7; blue circles), 200 (n=6; black circles) or 250–300 (n=7; red circles) post-pre pairings.Insets correspond to the average EPSC amplitude at baseline (black trace) and at 40–50 min after STDP pairings (grey trace). Error bars represent the SEM.

We next investigated whether a higher *N*_pairings_ would secure the expression of NMDAR-tLTP even with σ_Δ*t*_ >3 ms (Fig. 6B). When we doubled *N*_pairings_ we did not observe in average expression of significant tLTP. Indeed, 200 post-pre pairings (−30< *m*_Δ*t*_<0ms with 3<σ_Δ*t*_<10 ms) did not induce significant plasticity (112±9%, *p*=0.236, *n*=6; 2/6 cells displayed tLTP; Fig. 6B3); Nevertheless, it should be noted that with 200 pairings and with σ_Δ*t*_ >3 ms, tLTP were detected in two cases out of 6 trials whereas with 100 post-pairings a complete lack of tLTP was observed (n=7 cells). We thus increased the *N*_pairings_ up to 250–300. In these conditions, tLTP were reliably observed with σ_Δ*t*_ >3 ms. Figure 6B1 shows an example of tLTP induced by 250 pairings with σ_Δ*t*_ = 5 ms and centered on −20 ms where the mean baseline EPSC amplitude was 126±3pA before pairings and was increased by 57% to 198±4pA 45 minutes after pairings. Overall, 250-300 post-pre STDP pairings (−30< *m*_Δ*t*_<0 ms with 3< *m*_Δ*t*_<10 ms) induced tLTP (174±10%, *p*=0.0003, n=7; 7 cells out of 7 resulted in tLTP; Fig. 6B2 and B3); the mean values of these tLTP observed for *N*_pairings_ = 250–300 at 1 Hz with 4<σ_Δ*t*_<9 ms were not significantly differently than tLTP induced with 100 post-pre pairings at 1 Hz with σ_Δ*t*_<3 ms (*p*=0.1952).

### From spike timing- to frequency-dependent plasticity

We next tested with our mathematical model whether the robustness of STDP to jitter of the spike timing depends on the regularity of the spike train. In Figure 2E, we combined a triangular distribution for σ_Δ*t*_ and Poisson-distributed IPIs with a constant and relatively large refractory period *τ*_*r*_ (see *Methods*). However, when *τ*_*r*_ decreases, short IPIs are more frequently sampled and the train of presynaptic stimulation becomes more irregular. Therefore, through variations of the refractory period *τ*_r_, the presynaptic stimulation can be progressively switched from very irregular to very regular spike trains.

Figure 7 shows model predictions for the alteration of the robustness curves when *τ*_r_ varies. With high refractory period, ie. when τ_*r*_ ≈ *F*_pairings_, the pairings are very regular, so one recovers the robustness curves of Figure 2E, with eCB-STDP more robust than NMDAR-tLTP. However, when τ_*r*_ is 70% of *F*_pairings_, the robustness of both eCB-tLTP and NMDAR-tLTP increases and both exhibit similar robustness. With even smaller values of τ_*r*_, NMDAR-tLTP becomes more robust than eCB-tLTP. Moreover, the canonical eCB-tLTD (*F*_pairings_ = 1Hz, *m*_Δ*t*_ = +20 ms, *N*_pairings_=100) progressively stops triggering LTD when τ_*r*_ decreases, even inducing LTP for very low τ_*r*_. Hence, when the presynaptic spike train is progressively switched from very regular to very irregular spike trains, two main changes occur: (*i*) the three plasticity forms (NMDAR-tLTP, eCB-tLTD and eCB-tLTP) become increasingly robust to jittering. At the limit of very irregular spike trains, jittering hardly affects plasticity amplitudes; (*ii*) eCB-tLTD is progressively changed to LTP, so that very irregular paired stimulations with *F*_pairings_=1Hz only produce LTP regardless the sign of the spike timing.

**Figure 7:**
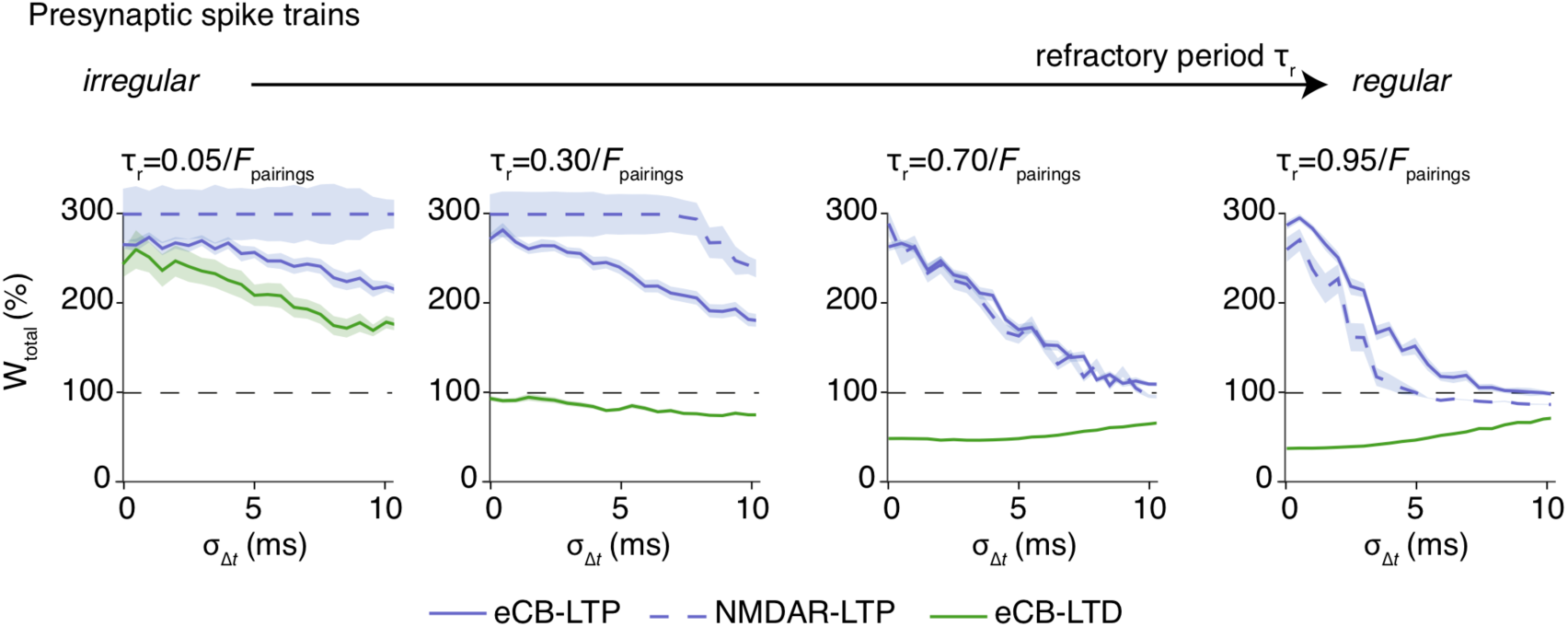
Plasticity robustness depends on the regularity of the presynaptic spike train. The figure shows the robustness of the model plasticity (eCB-tLTP with purple, eCB-tLTD with green and NMDAR-tLTP with dashed purple) with the jittered STDP protocol of figure 2E1, i.e. exponentially-distributed IPIs with refractory period r_r_ and triangularly-distributed jitter *ξ*_Δ*t*_. From the right-hand side to the left-hand side of the figure, the refractory period τ_*r*_ is gradually decreased. When τ_*r*_ → 1/F_pairings_ where F_pairings_ = 1 Hz is the average pairings frequency, one gets regular presynaptic spike trains, similar to classical STDP protocols. In this case (see e.g. τ_*r*_ = 0.95/F_pairings_), the robustness pattern is similar to the previous observations at 1 Hz: eCB-STDP is very robust, whereas NMDAR-tLTP is fragile. However, when τ_*r*_ → 0, the presynaptic spike train becomes very irregular. In this case, the robustness patterns changes totally, with NMDAR-LTP becoming the most robust, and eCB-tLTD turning into a LTP. Hence, irregular presynaptic spike trains are predicted to produce very robust LTP at the expense of LTD.

## DISCUSSION

In the present study, we have introduced jittered STDP protocols as noisy variants of STDP protocols. In jittered STDP, both the spike timing, i.e. the delay between a pre- and paired postsynaptic stimulation (or vice-versa), and the pairing frequency become random variables. Our mathematical model of the signaling pathways that underlie STDP predicted that the expression of STDP is robust to some amount of jittering of the spike timing. However, STDP eventually disappears in the model when the amplitude of the jitter becomes very large i.e. when its standard deviation σ_Δ*t*_ > 3 ms for NMDAR-mediated plasticity and σ_Δ*t*_ > 10 ms for eCB-mediated plasticity. We tested this prediction experimentally by single neurons recordings with patch-clamp and by applying jittered STDP to corticostriatal synapses between somatosensory cortex and the dorso-lateral striatum. Indeed, at these synapses it exists a variety of STDP forms, that involve at least two signaling pathways, NMDAR and endocannabinoids [8,9,15,16,17,19]. Our experiments confirmed the predictions of the model, both qualitatively and quantitatively. Therefore, in a noisy neural network, STDP is expected to resist low to moderate variability but to vanish upon higher noise. However, the magnitude of variability that STDP can tolerate varies strongly depending on the pairing frequency and on the STDP form, i.e. NMDAR-tLTP, eCB-tLTP or eCB-tLTD.

At 1 Hz pairing frequency, we show that eCB-STDP (eCB-tLTD and eCB-tLTP) is from far more robust than NMDAR-dependent tLTP. However, we found that this robustness depends on the number and frequency of pairings. Indeed, increasing the average pairing frequency to 3 Hz improved the robustness of NMDAR-tLTP. We observed a similar improvement of the robustness of NMDAR-tLTP to jitter when the number of pairings increased more than two folds. These results have wide-ranging implications for STDP expression in *in vivo*-like firing. They indicate that eCB-STDP could be more likely responsible for fast learning involving very few trails (eCB-tLTP) or learning at low activity frequency than NMDAR-STDP. NMDAR-STDP could however take part in learning for larger frequencies or when the same spike timing recurs a large number of times during sustained activity. It also implies that in synapses where NMDAR is the main coincident detector for tLTD or tLTP [17,29,30,31,32,33,34], a high degree of precision of the spike timing, and/or an increased number or frequency of pairings, are required for the emergence of the NMDAR-mediated plasticity. This can be viewed as a mechanism preventing the occurrence of spurious plasticity for too noisy neural network activity. At synapses in which NMDAR and endocannabinoids are both involved in STDP expression [8,9,16,35,36,37,38] and in which eCB-LTP can occur, as it is the case in the striatum [8,9] or the hippocampus [39,40], the emergence of STDP would be possible even in noisy conditions and could thus serve subsequently as a priming for the subsequent expression of NMDAR-tLTP.

Another contribution of the present study is the exploration of the robustness of STDP when noise affects not only the spike timing but also the inter-pairing interval (IPI), i.e. the delay between two successive pairings. In the absence of jitter of the spike timing, our model predicts that IPI irregularity tends to consolidate tLTP at the expense of tLTD. This prediction that irregular IPIs strengthen tLTP when the spike timing is constant confirms the result obtained independently and with different mathematical models (see Fig. 3 in [13]). When IPIs are regular, adding jitter of the spike timing in our model progressively suppresses STDP. However, with more irregular IPIs our model predicts that the three STDP forms should exhibit larger robustness to noise. Therefore our result suggests that jittering of the spike timing has larger impact on STDP when the IPIs are regular, but that jittering has much less consequence with very irregular IPIs. Experimental testing of this model prediction would greatly improve our understanding of the induction and maintenance of STDP in *in vivo-*like firing.

Mathematical models in neuroscience can have two, very different, purposes [41]. One purpose can be to gain a theoretical account of the studied experimental system, i.e. a synthetic and formal understanding of the system’s behavior, usually related to how it transfers or computes information. However, this is not always of practical value for experimental neurobiologists because it is difficult to transpose pharmacological experiments in these models since they abstract out the details of the molecular levels. Here, we used another type of models that consists in describing individually all the important molecular reactions using biophysical laws or biochemical kinetics. Such systems-biology-like models are too complex to allow the level of first-principle understanding provided by theoretical models but they have two main advantages: (*i*) pharmacology experiments are very easily mapped in the model and (*ii*) if the model behavior is similar enough to that of the experimental system, the model can be used to select, among the vast number of possible experiments, what experiments are the more useful to validate or reject hypothesizes. For instance, in electrophysiology, it is hardly possible to fully explore every possible parameter ranges of STDP. Therefore, in the experiments we have fixed the distribution function of the spike timing jitter to a Normal distribution (i.e. Fig. 2C1). We have then used our model to explore other distributions (uniform, triangular, with or without jitter of the interval between pairings) and consistently obtained the same results for all these distributions. This strongly suggests that our main result, *i.e*. the differential robustness of STDP forms to jitter, with eCB-STDP being more robust than NMDAR-tLTP, does not depend on the distribution function of the jitter.

The existence of synaptic plasticity induced by a STDP-like protocol without postsynaptic spikes has been reported in the striatum [42] and hippocampus [43]. In addition, in the hippocampus, a low-frequency stimulation at 1 Hz induced LTD regardless of the magnitude of postsynaptic depolarization (i.e. sub- vs suprathreshold) [44]. It thus appears that the subthreshold postsynaptic depolarization is a key factor in the induction of plasticity [45]. In the striatum, we have previously described the existence of a potent plasticity induced by a subthreshold postsynaptic depolarization in the absence of postsynaptic spike (subthreshold depolarization-dependent plasticity, SDDP) [42]. However, this SDDP differs from STDP on two aspects: (*i*) SDDP was induced in a larger temporal window (−100< Δ*t*_STDP_<+100 ms) than STDP (−30< Δ*t*_STDP_<+30 ms) and (*ii*) in SDDP, tLTP and tLTD were indifferently induced regardless of the spike timing whereas in STDP Δ*t*_STDP_ appears as a key determinant of the orientation of the plasticity. Therefore, the action potential does not seem to be necessary to induce plasticity at the corticostriatal synapse, but would be determinant for the polarity and bidirectional characteristic of STDP and the width of the Δ*t*_STDP_ window. The effects of timing jitter on SDDP remain to be determined. Our model could be used to bring elements of answer to this question since it is based on the signaling pathways implicated and can accommodate any form of postsynaptic stimulation. It could therefore directly be used to test SDDP-like stimulation protocols in the presence of jitter.

The present paper focuses on the corticostrial synapse and its associated anti-Hebbian STDP, where tLTP is obtained with negative (post-pre) pairings whereas positive (pre-post) pairings give rise to tLTD. At corticostriatal synapses, we have shown that GABA controls the polarity of STDP [17,18,27], meaning that Hebbian [15,16] or anti-Hebbian [9,19,26,46] STDP were observed depending on whether GABA_A_Rs antagonists were applied [18,27]. In the cortex or the hippocampus, STDP is most often Hebbian: tLTP is expressed for positive spike timings and tLTD for negative ones [3,4,14]. Despite the fact that the shape of the dependence of STDP on (regular) spike timing presents a striking variety [5], the underlying signaling pathways are conserved [4,14]. One interpretation of the existence of this variety given the conservation of the underlying signaling pathways is that modulating the expression levels of (part of) the signaling molecules of those pathways may be enough to change the shape of STDP. For instance, modulating GABA signaling can switch STDP from Hebbian to anti-Hebbian in the striatum [18,27]. Likewise, modulating glutamate uptake has very strong effects on the shape of STDP [46]. The shape of the dependence of STDP on spike timing is also strongly altered by neuromodulatory signaling pathways, including dopamine [15,16,26,47,48], acetylcholine [49,50,51] and noradrenaline [48,51,52]. A major conclusion from our present work is that the robustness of STDP to jitter strongly depends on the underlying signaling pathways. For instance, the robustness of NMDAR-tLTP in our model is strongly dependent on the amplitude of the activated CaMKII transients triggered by each post-pre pairings or on alterations of the ratio between the decay time of these transients and the IPIs. The robustness of STDP to jitter could be similarly controlled by quantitative variations in the underlying pathways. Such variations are expected to occur between two neuron subtypes or brain regions but also as a result of the activation of a neuromodulatory pathway. Therefore, our work suggests that the expression of STDP in *in vivo*-like firing might appear or disappear as a result of modulations of its robustness to jitter, depending on the properties of the incoming patterns (reflecting for example fast learning or heavy training) but also the brain region and, more transiently, on the activation of neuromodulatory pathways.

